# Causal roles of frontoparietal cortical areas in feedback control of the limb

**DOI:** 10.1101/2020.05.02.072538

**Authors:** Tomohiko Takei, Stephen G. Lomber, Douglas J. Cook, Stephen H. Scott

## Abstract

Goal-directed motor corrections are surprisingly fast and complex, but little is known on how they are generated by the central nervous system. Here we show that temporary cooling of dorsal premotor cortex (PMd) or parietal area 5 (A5) in behaving monkeys caused impairments in corrective responses to mechanical perturbations of the forelimb. Deactivation of PMd impaired both spatial accuracy and response speed, whereas deactivation of A5 impaired spatial accuracy, but not response speed. Simulations based on optimal feedback control demonstrated that ‘deactivation’ of the control policy (reduction of feedback gain) impaired both spatial accuracy and response speed, whereas ‘deactivation’ in state estimation (reduction of Kalman gain) impaired spatial accuracy but not response speed, paralleling the impairments observed from deactivation of PMd and A5, respectively. Furthermore, combined deactivation of both cortical regions led to additive impairments of individual deactivations, whereas reducing the amount of cooling (i.e. milder cooling) to PMd led to impairments in response speed, but not spatial accuracy, both also predicted by the model simulations. These results provide causal support that higher order motor and somatosensory regions beyond primary somatosensory and primary motor cortex are involved in generating goal-directed motor responses. As well, the computational models suggest that the distinct patterns of impairments associated with these cortical regions reflect their unique functional roles in goal-directed feedback control.

## Main text

The motor system can generate a broad range of motor actions and flexibly adjust them to manage errors or unexpected changes in the environment. It is commonly assumed that these goal-directed motor corrections are generated through a rapid transcortical pathway involving primary somatosensory (S1) and primary motor cortex (M1), leading to muscle responses in as little as 60ms for the arm^1–3^. However, our recent study highlighted that short latency neural responses to mechanical disturbances applied to the forelimb of a monkey can be observed in as little as 25ms across frontoparietal circuits, including in higher order motor (dorsal premotor cortex, PMd) and somatosensory regions (parietal area 5, A5)^4^. These cortical regions are normally associated with motor planning and movement initiation^5,6^, yet their contribution to online motor control remain poorly understood. The objective of this study is to provide causal support that these cortical regions are also involved in goal-directed feedback control by quantifying how transient deactivation of each cortical region, induced with cortical cooling^7^, impacts goal-directed feedback responses.

How could changes in a feedback circuit alter goal-directed feedback responses? We made theoretical predictions for the motor deficits that may occur from deactivating a brain region using an optimal feedback control (OFC) model, a common framework for interpreting voluntary motor function (Fig. 1a)^8,9^. We first optimized model parameters to reproduce the monkey’s intact feedback response (Fig. 1b). Then, we applied ‘deactivation’ to the model parameters (Fig. 1a), including (1) the feedback gain of the control policy (*L*), (2) Kalman gain in state estimation (*K*), (3) parameters in the internal forward model (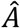, 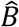 and 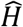), and (4) the sensory observation matrix (*H*). Since cooling reduces neural excitability in adjacent tissue^7^, we modeled a cooling effect as a reduction of each parameter (for a different deactivation method based on increases in noise, see supplementary discussion). First, large reductions in any parameter ultimately led to failure of the model to generate appropriate motor corrections to mechanical disturbances (Fig. 1d). Of particular interest is that smaller reductions of parameters led to unique patterns of feedback impairments for each parameter (Fig. 1c-d). Reduction of the feedback gain (*L*), that converts estimates of system state into motor commands, generates a broad range of impairments, including reduction in spatial accuracy (endpoint error) and several metrics related to response speed (return time, max deviation and max deviation time, Fig. 1c-d, red, *p* < 0.05). Reduction of Kalman gain (*K*) degrades the use of external sensory feedback, making state estimation more reliant on internal feedback. This generated impairments in spatial accuracy (endpoint error), but interestingly, did not impair response speed (return time and max deviation time) until the gain was reduced more than ~60% (Fig. 1c-d, blue, *p* > 0.05). This reflects that reduction of the Kalman gain induces only a small delay in updating the state estimation about the presence of a disturbance using external sensory feedback, but once updated, internal feedback can then appropriately counter the external perturbation. Even small reductions of parameters in the internal forward model (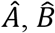 and 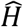) providing internal feedback for state estimation quickly leads to severe oscillations (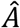, Fig. 1c, green; 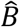 and 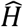, Fig. S1a), with rapid degradation in spatial accuracy (Fig. 1d, green and Fig. S1b, *p* < 0.05). Finally, reduction of the sensory observation matrix (*H*) degrades sensory information. This generated impairments in spatial accuracy and response speed (Fig. 1c-d, orange, *p* < 0.05), qualitatively similar to impairments associated with the feedback gain (*L*). Taken together, our simulations highlight that deactivation of different model parameters generate unique patterns, or signatures, of motor impairments which can be used to dissociate cortical functions.

**Figure 1.**
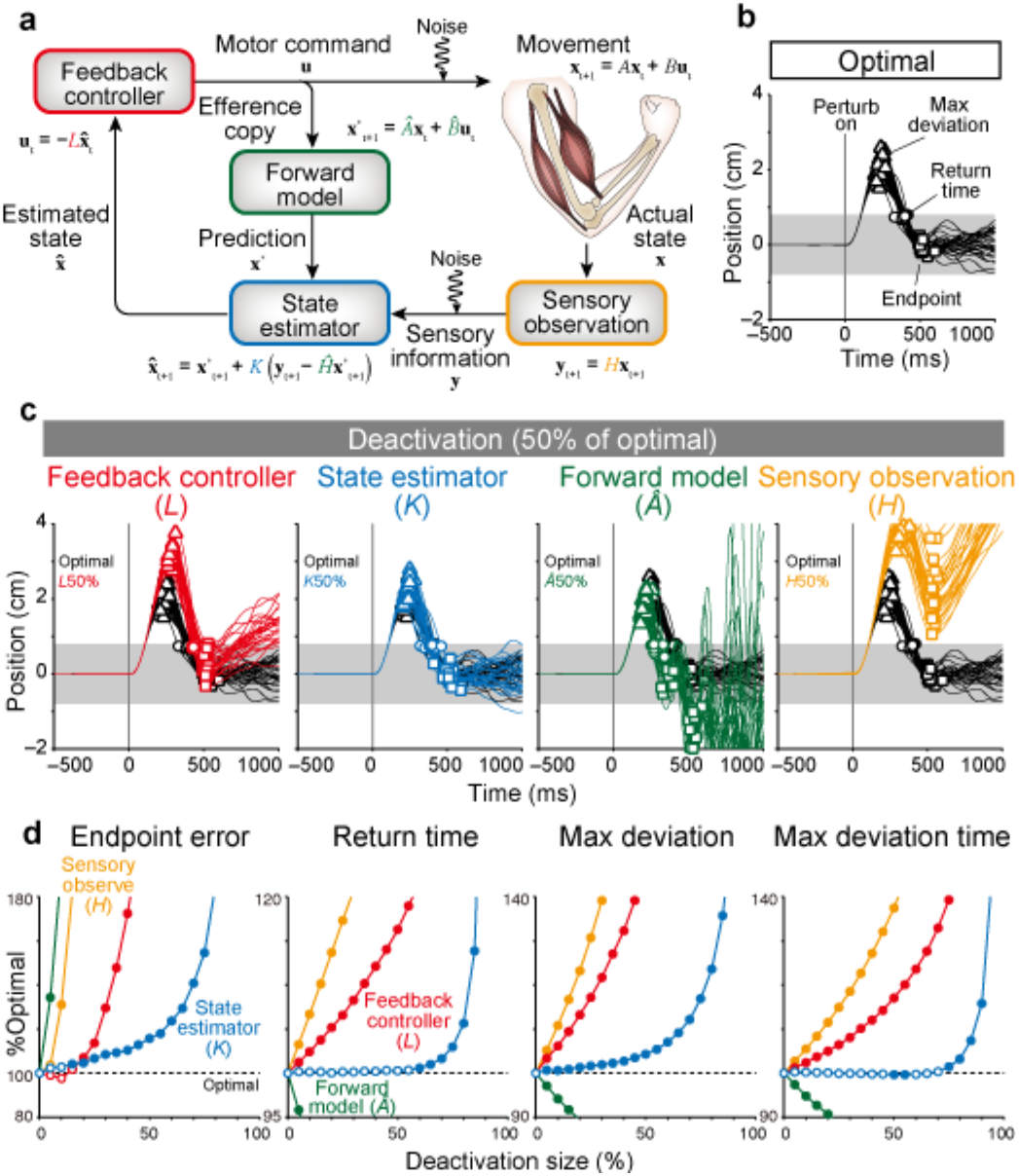
Simulation for the effect of cortical deactivations on feedback motor responses to mechanical perturbations. **a**, Optimal feedback control model. **b,** Response of the model to mechanical perturbations in optimized condition. **c**, Response of the model in deactivated conditions. Black lines denote the optimized condition (same to **b**). Coloured lines denote trials when each parameter was reduced by 50%. Red, feedback gain in feedback controller (*L*). Blue, Kalman gain in state estimator (*K*). Green, a parameter of forward model 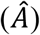. Yellow, observation matrix in sensory observation (*H*). **d**, Performance measures impaired by the deactivation of model parameters. All measures showed significant interaction and main effects in two-way ANOVA (deactivation size × deactivated parameters) suggesting unique patterns of motor impairments for each parameter (*p* < 0.05). Filled circles, significant difference from optimal condition (*t*-test, *p* < 0.05).

The traditional view is that frontal motor cortex is involved in ‘motor’ processing and parietal cortex is involved in ‘sensory’ processing^5,6^. Thus, we hypothesized that PMd cooling would parallel impairments associated with reductions in the feedback gain (spatial accuracy and response times) and A5 cooling would parallel impairments associated with the Kalman gain (only spatial accuracy).

To test our hypotheses, we chronically implanted cooling probes over the surface of PMd and A5 (Fig. 2a). By circulating chilled methanol through the probes, we are able to cool down the cortical temperature and deactivate each area^7^ while observing changes in motor performance during a single experimental session reversibly and temporarily (Fig. 2b). Monkeys were trained to maintain their fingertip at a central target and to make corrections to mechanical perturbations unexpectedly applied to the forelimb (posture perturbation task^10^, Fig. 2c). On separate days, the task was performed before (pre-cool), during (cool) and after cooling (post-cool) each cortical region, including sham controls when cooling was not applied (monkey A: PMd n=23, A5 n=31, Sham n=28; monkey R: PMd n=10, A5 n=18, Sham n=51 sessions). When PMd was cooled, motor responses were slowed (Fig. 3a, bottom) and response accuracy was reduced (Fig. 3a, top), resulting in a significant increase in endpoint errors (*t*(79.7) = 4.8, *p* < 0.05) and response speed (Fig. 3c, return time, max deviation and max deviation time, *t*(87.2) = 5.7, *t*(83.8) = 4.4, *t*(88.1) = 5.4, *p* < 0.05, respectively). Correspondingly, the muscle stretch response was reduced beginning in the long-latency time epoch (R3 epoch, 75-120ms after perturbation onset, *t*(92.3) = 3.6, *p* < 0.05, Fig. 3d), which is the first instance that transcortical feedback can contribute to feedback corrections^1,2,11,12^. These results are consistent with our hypothesis that PMd is involved in the feedback control policy. We confirmed that these behavioural effects were consistent between two monkeys (Fig. S2) and not due to direct cooling of M1 (Fig. S3).

**Figure 2.**
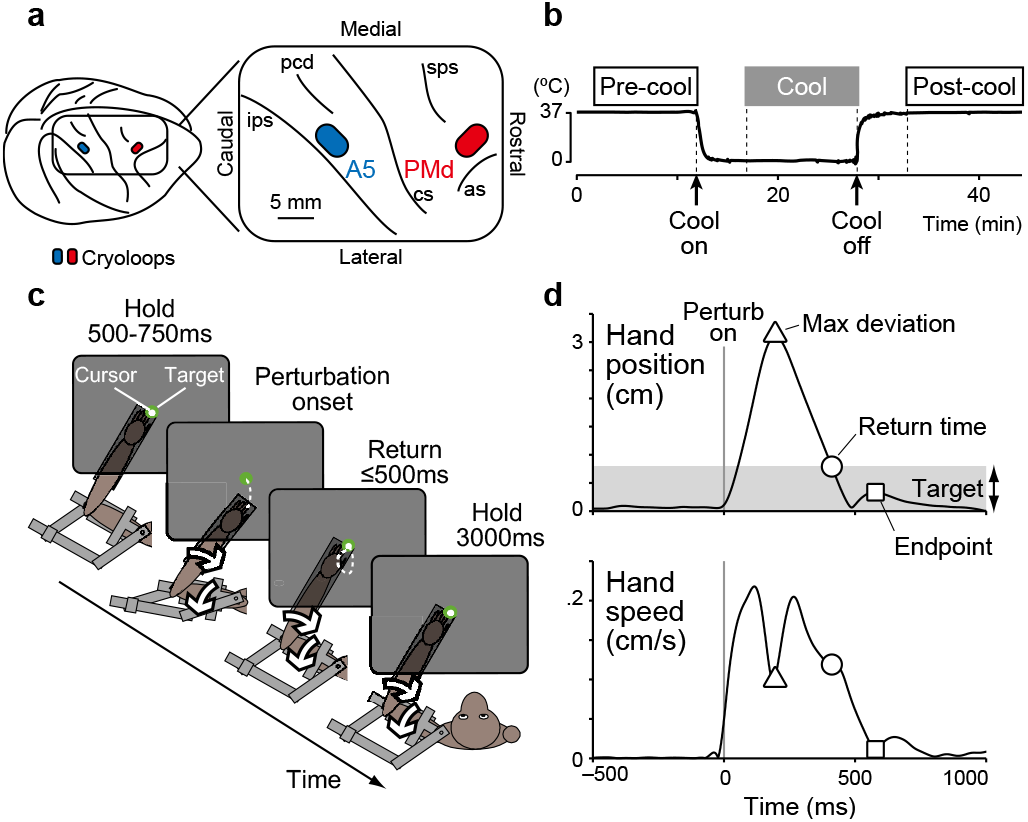
Experimental setups for cortical cooling. **a**, Cooling probes were implanted over dorsal premotor cortex (PMd) and parietal area 5 (A5) in the right hemisphere of monkeys. **b**, Feedback motor responses were tested before (pre-cool), during (cool) and after (post-cool) cooling. **c**, Postural perturbation task. Monkey must maintain its hand at a central target. A mechanical load was applied to the limb and the monkey must return its hand to the same spatial target in less than 500 ms and maintain its hand for another 3 seconds. **d**, Hand position and speed in an exemplar trial. Symbols denote behavioural measures: max deviation (triangle), return time (circle) and endpoint (square).

**Figure 3.**
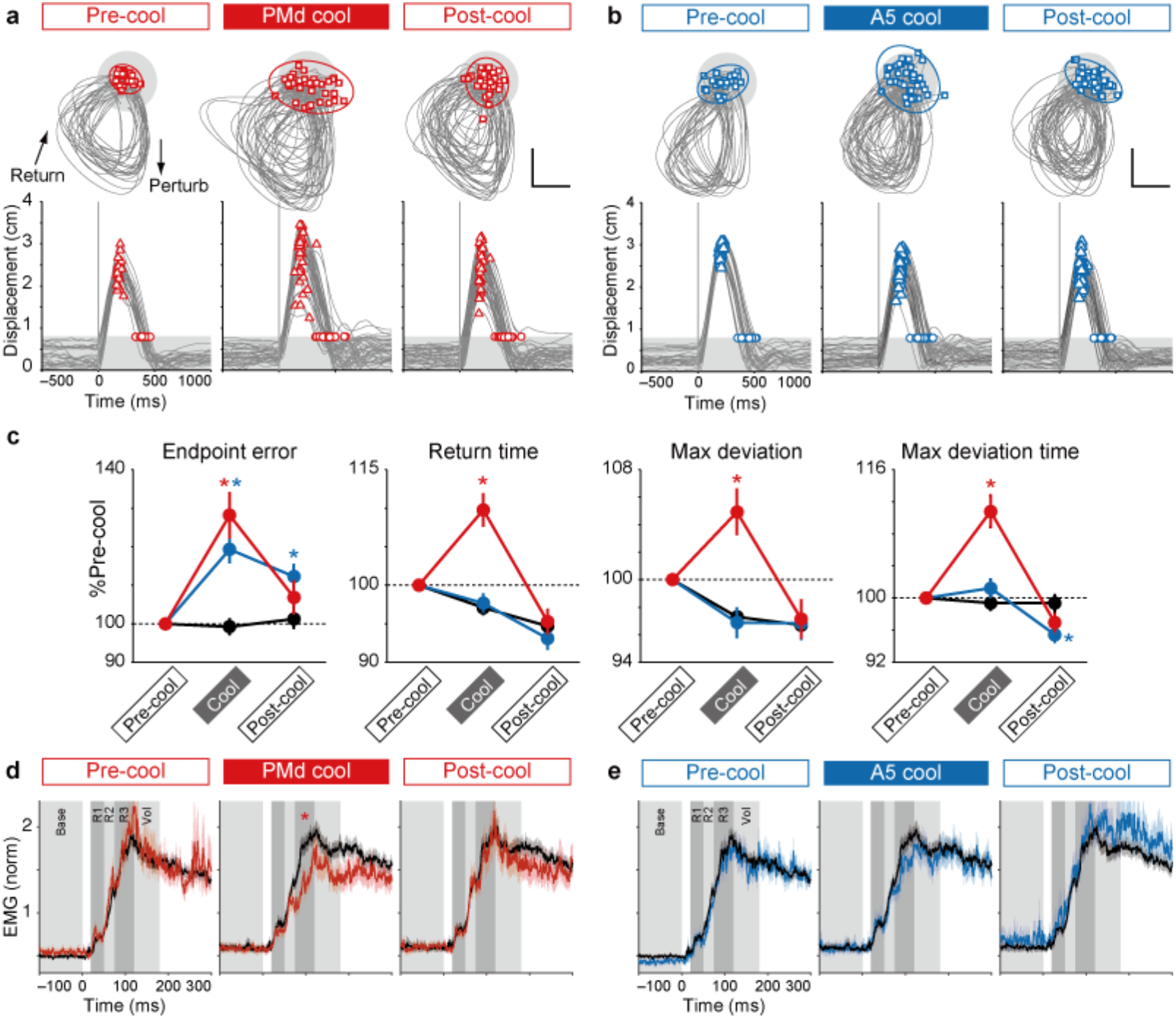
Effects of PMd and A5 cooling. **a**, **b**, Hand trajectories (top) and hand displacements (bottom) when the limb was unexpectedly perturbed in pre-cool, cool and post-cool PMd conditions (**a**) and corresponding A5 conditions (**b**). Calibration bars, 1cm. Ellipses, 95% confident interval of endpoints. Time = 0, perturbation onset. **c**, performance measures for PMd (red), A5 (blue) and sham (black) cooling sessions. Averages across two monkeys and two torque directions after normalized to pre-cool condition. Error bars, SEM. All measures displayed significant interaction and main effects in two-way ANOVA (cooling epochs × target areas) indicating different effects of cortical cooling (*p* < 0.05). * Significant difference from sham condition (*t*-test, *p* < 0.05). **d**, **e**, Averaged EMG responses to mechanical perturbations in PMd (red), A5 (blue) and sham (black) cooling sessions. Responses were binned to baseline (100-0 ms before perturbation), R1 (20-50 ms post-perturbation), R2 (50-75 ms), R3 (75-120 ms), and voluntary (120-180 ms) time windows (gray and dark gray rectangles). Shaded areas, SEM. * Significant difference from sham condition (*t*-test, *p* < 0.05).

In contrast, A5 cooling significantly increased endpoint errors (*t*(162) = 5.3, *p* < 0.05), but did not impair response speed (return time, max deviation and max deviation time, *t*(200) = 0.4, *t*(162) = 0.4, *t*(201) = 1.4, *p* > 0.5, respectively), suggesting that the earlier motor response was preserved (Fig. 3b-c), which was confirmed by the lack of significant changes in EMG during the R3 epoch (*t*(63) = 1.8, *p* = 0.42, Fig. 3d). These results are consistent with our hypothesis that A5 is involved in state estimation.

Since our simulation showed that reductions of feedback control gain (*L*) and sensory observation (*H*) caused qualitatively similar effects (Fig. 1d), we could not separate whether PMd cooling impaired feedback gain or sensory observation. Therefore, we further dissociate their effects by simulating the simultaneous cooling of PMd and A5 with a simultaneous reduction of *L* and *K* (*L* & *K*), or *H* and *K* (*H* & *K*). We found that reduction of *L* & *K* led to impairments that were a linear sum or a sublinear interaction of impairments for deactivation of each parameter separately (Fig. S4a). In contrast, reduction of *H* & *K* induced supralinear impairments of each parameter separately (Fig. S4b). When we simultaneously cooled PMd and A5 (n = 20 and 10 in monkey A and R), we found that impairments were a linear sum (endpoint error, return time and max deviation time) or a trend of the sublinear interaction (max deviation) of impairments induced by individual cooling of each area separately (Fig. 4a, endpoint error, return time, max deviation time, *t*(59) = 0.7, *t*(59) = 0.1, *t*(59) = 0.3, *p* > 0.9; max deviation,, *t*(59) = 1.2, p = 0.47). These results provide further support for our hypotheses that PMd and A5 are involved with the feedback control policy (*L*) and state estimation (*K*), respectively.

**Figure 4.**
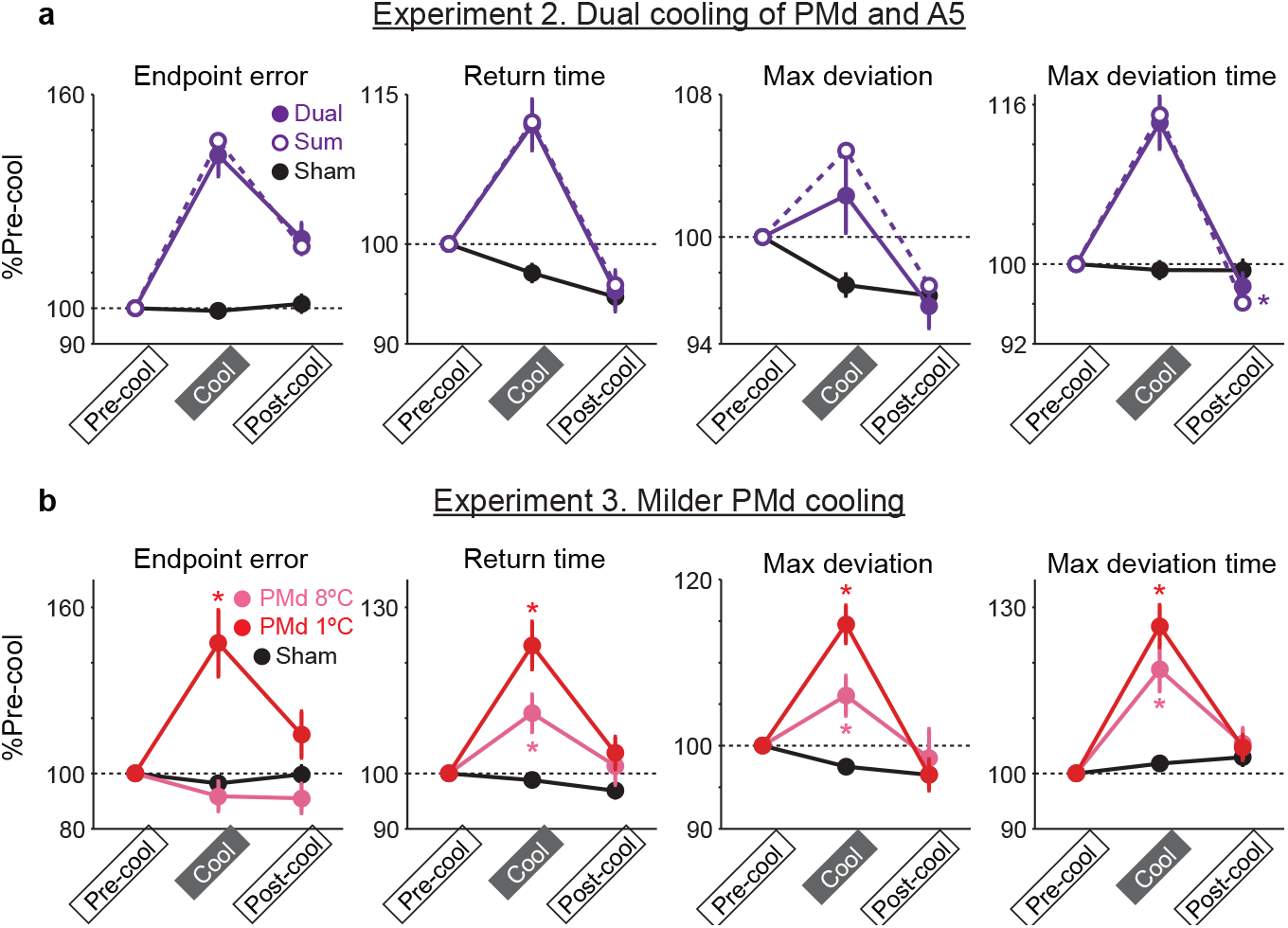
Effects of dual cooling and milder PMd cooling. **a**, Performance measures of dual cooling of PMd and A5 (filled circle, “Dual”) were compared with a linear summation of the effects of single PMd and A5 cooling (open circle, “Sum”). * Significant difference between dual cooling and linear sum (*t*-test, *p* < 0.05). **b**, Comparison of original cooling (1°C in probe temperature, red) and milder cooling of PMd (8°C, pink). * Significant difference from sham condition (*t*-test, *p* < 0.05). Error bars, SEM. All measures showed significant interaction and main effects in two-way ANOVA (cooling epochs × target areas) indicating different effects of cortical cooling (*p* < 0.05).

Finally, we tested a further prediction of the model that a small reduction in the feedback gain *L* (≤ 20%) would lead to a unique pattern of impairments in which most motor parameters would be affected (return time, max deviation, and max deviation time) but not endpoint error (e.g. 10% deactivation of *L* in Fig. 1d). We tested this prediction by applying milder cooling of PMd in one animal (8 °C instead of 1 °C in probe temperature, n = 5 in monkey R). Results showed that less cooling significantly impaired all parameters (return time, max deviation, max deviation time, *t*(11) = 3.2, *t*(10) = 3.3, *t*(10) = 4.0, *p* < 0.05, respectively) except endpoint errors (Fig. 4b, pink, *t*(12) = 0.8, *p* = 1.0). This result further validates our simulation model of cortical cooling.

Taken together, these results provide clear support that PMd and A5 are causally involved in generating goal-directed feedback responses. Interpreting their potential contribution to feedback responses was possible by comparing the pattern of impairments associated with deactivation of each cortical region with simulation of deactivation using an OFC model. We expect qualitatively similar results would be observed for other control models that possess a control policy and state estimation that integrates internal feedback with external sensory feedback, known features of the voluntary motor system^13,14^. It is interesting to note that reductions in parameters associated with the forward model led to oscillatory behavior (Fig. 1c and Fig. S1a), which resembles ataxia of cerebellar patients and a pattern of impairment when the dentate nucleus in the cerebellum was cooled^15,16^. Comparison of signatures of motor impairments of cortical deactivations and model simulations can be a powerful tool to unravel the computational basis of sensorimotor systems and their dysfunctions.

Monkeys were still able to generate goal-directed responses during cortical cooling, likely reflecting that only a portion of these cortical regions were impacted by the cooling and/or that these processes are distributed across brain regions^3,17^. For example, M1 is almost certainly involved in processes associated with the control policy^1–3,11,18–20^. It is possible that deactivation of PMd, traditionally associated with motor planning, caused motor impairments by degrading ongoing input to M1^21–23^. However, PMd may also have a more direct influence in generating motor corrections through its corticospinal projections^24^. Recent studies highlight how neural activity can reflect multiple distinct processes using orthogonal subspaces^25–29^. For example, M1 can reflect simultaneous motor actions of different limbs with the associated neural activity for each limb segregated to orthogonal subspaces^29^. Thus, it is plausible that PMd could simultaneously be involved in motor planning in one subspace, while it also contributed to online feedback control in another orthogonal subspace.

## Acknowledgements

We thank Kim Moore, Simone Appaqaq, Justin Peterson, Helen Bretzke and Mike Lewis for their laboratory and technical assistance. We also thank Dr. Mohsen Omrani for his support to setup the cooling system. This work was supported by grant support from the Canadian Institutes of Health Research (CIHR) and The Uehara Memorial Foundation, The Naito Foundation, The Takeda Science Foundation and Japan Society for the Promotion of Science Grants-in-Aid (19H03975, 19H05311).

## Author Contributions

Conceptualization, T.T. and S.H.S.; Methodology, T.T., S.G.L., D.J.C. and S.H.S.; Investigation, T.T.; Writing – Original Draft, T.T. and S.H.S.; Writing – Review & Editing, T.T., S.G.L., D.J.C. and S.H.S.; Funding Acquisition, S.H.S; Supervision, S.H.S.

## Author information

S.H.S. is co-founder and CSO of Kinarm Technologies that commercializes the robotic technology used in this study.

Correspondence and requests for materials should be addressed to T.T. or S.H.S.

## Methods

### Subjects and apparatus

Two male rhesus monkeys (Macaca mulatta, 10–17 kg, monkeys A and R) were used in this study following procedures approved by the Queen’s University Animal Care Committee. They were trained to perform upper limb motor tasks with their left arm while wearing a robotic upper-limb exoskeleton (Kinarm Exoskeleton; Kinarm Technologies, Kingston, Ontario, Canada) that permitted and monitored horizontal shoulder and elbow motion^30^. A virtual reality system presented visual targets and a cursor representing hand position in the workspace while direct view of their limb was occluded.

### Behavioral task

We trained monkeys to perform a posture perturbation task, which has been described previously^10^. In the task, the monkey was required to maintain a small cursor (0.2 cm radius) representing the position of the index fingertip at a visual target (0.6 cm radius) displayed near the center of the arm’s workspace (~30° and 90° degrees at the shoulder and elbow joints, respectively, Fig. 2c). The monkeys initiated each trial by moving their hand to the visual target and maintaining it within the target’s acceptance window for 0.5 to 2.5 s (monkey A) or from 0.5 to 1.25 s (monkey R). The size of the acceptance window was individually adjusted to each monkey (0.8 and 1.0 cm radius for monkey A and R, respectively). Then, one of three mechanical loads was applied to the monkeys’ arm. The load conditions included shoulder extension and elbow flexion (SE+EF), shoulder flexion and elbow extension (SF+EE), and an unloaded condition (catch trials). These loads stayed until the end of the trial (a step-torque perturbation) and the monkeys were required to counter the load to return to the target within 0.5 s and maintain it there for another 3.0 s (monkey A) or 2.0 s (monkey R) to receive a liquid reward. Shoulder and elbow torques of magnitude 0.28 Nm (monkey A) or 0.20 Nm (monkey R) were used. Each block consisted of four SE+EF trials, four SF+EE trials and one catch trial in random order (total nine trials per block). To mitigate the contribution of visual feedback for the rapid motor responses, we removed the hand cursor feedback for 200 ms after perturbation onset.

### Cortical Cooling

After behavioural training was complete, we performed surgical procedures to implant two cooling probes over PMd and A5 as well as a head fixation post. In monkey A, we also made a craniotomy over M1 and implanted a recording chamber to record the temperature of M1. The surgeries were performed using isoflurane anesthesia (1.0–2.0% in O2) and under aseptic conditions. PMd and A5 locations were identified based on sulcus structures on the cortical surface according to a previous electrophysiological study^31^. PMd probe was implanted between upper limb of arcuate sulcus and superior precentral sulcus, whereas A5 probe was implanted between intraparietal sulcus and postcentral dimple (Fig. 2a).

Cooling probes were made with 23G stainless-steel tubing in 3 × 5 mm dimension^7^. A thermocouple is attached to the tip of the probe to monitor probe temperature. By circulating chilled methanol, the probe temperature was controlled at the desired temperature. Previous work has shown that cooling the probe to 1°C deactivates cortex up to a distance of 1.5 mm, which covers most of the cortical layers^7^. Thus, the estimated volume of deactivated cortical tissue is 72–126 mm^3^.

### Experimental procedure

On each experimental day, one of the cooling conditions was chosen. In experiment 1, one of the probes (PMd and A5) was cooled or no cooling was applied (sham). In experiment 2, PMd and A5 was simultaneously cooled (dual cooling). In experiment 3, PMd in monkey R was cooled to a milder target temperature (8 ± 1°C in probe temperature) instead of 0 – 1°C (milder cooling).

Each experiment was initiated with a brief practice set (3 and 2 blocks of trials for monkey A and R, respectively) with a normal acceptance window (0.8 and 1.0 cm radius for monkey A and R). After that, the pre-cooling epoch started (Fig. 2b). In the pre-cooling epoch, the monkeys performed 6 (monkey A) or 3 blocks (monkey R) of trials, which consisted of 48 or 24 perturbed trials and 6 or 3 catch trials, respectively. Since we expected a motor impairment during cooling, we relaxed the acceptance window to 1.8 and 2.0 cm radius (monkey A and R) and we used the same window during all epochs (pre-cool, cool and post-cool) except for the initial practice set.

After the pre-cooling epoch, we started to circulate chilled methanol through the cooling probes and manually controlled the flow to keep the probe temperatures at the target temperature (0 – 1°C for experiment 1 and 2 and 8 ± 1°C for experiment 3). It took 80 ± 39 sec to reach the target temperature (0 – 1°C). After that, we waited ~5 min for the temperatures to become stable. Then we collected behavioural data in the cooling epoch. In the cooling epoch, the monkeys performed 9 (monkey A) or 5 – 6 blocks (monkey R), which consisted of 72 or 40 – 48 perturbed trials and 9 or 5 – 6 catch trials, respectively. It took ~20 min to complete all blocks.

Then, methanol circulation was stopped and after ~5 min the post-cooling epoch started. It took 126 ± 45 sec for the temperature to return back to normal (> 36°C). In the post-cooling epoch, the monkeys performed 9 (monkey A) or 5 – 6 blocks (monkey R), which consisted of 72 or 40 – 48 perturbed trials and 9 or 5 – 6 catch trials, respectively. In some recording sessions (9/94 sessions), monkey R did not complete the post-cooling epoch and we included only pre-cool and cool epoch data. We verified that inclusion of these incomplete sessions did not qualitatively affect the cooling results.

### Data analyses

All subsequent analyses were performed off-line using MATLAB (The MathWorks, RRID: SCR_001622).

### Kinematics analyses

Kinematic data and applied torques were acquired directly by the Kinarm device and were sampled at 4000 Hz with Plexon systems (Plexon Inc., Dallas TX, USA) and low-pass filtered (6th order double-pass filter, cutoff = 10Hz). We quantified four behavioural measurements related to spatial accuracy and response speed of the mechanical perturbation (Fig. 2d).

Endpoint error was quantified to measure spatial accuracy of the feedback response. First, we identified the endpoint of the initial corrective response after the perturbation. To do this, we calculated radial hand velocity relative to the center of the target and identified the timing of the maximum inward (i.e. returning) velocity. After this time point, we sought the first timing when the monkey’s hand had stopped moving according to a two-threshold method^32^: (1) the first local minimum in hand speed below 0.05m/s or (2) when hand speed dropped below 0.005m/s (endpoint, Fig. 2d square). Endpoint error was measured as a distance between hand position at the endpoint and the center of the target.

Return time was measured as the time interval from perturbation onset to when the hand cursor reentered the target area. Practically, we used a larger acceptance window (1.8 and 2.0 cm for monkey A and R) to keep animals rewarded even with motor impairments. However, when we calculated the return time, we used the original target size (0.8 and 1.0 cm for monkey A and R) that the monkeys were trained with (Fig. 2d circle).

Max deviation was defined as the peak hand displacement from the pre-perturbation hand position (Fig. 2d triangle). Pre-perturbation hand position was calculated as the averaged hand position during an interval from 100 to 0 ms before perturbation onset. Max deviation time is the time when max deviation occurred relative to perturbation onset.

Modulation of behavioural measures was tested by using a two-way ANOVA (cooling epochs × target areas, *p* < 0.05). As *post hoc* analyses, Welch’s *t*-tests were performed to compare between cooling and sham conditions (*p* < 0.05 with Bonferroni correction). Since we verified the difference of torque directions (SE+EF or SF+EE) did not qualitatively affect cooling results, we pooled the data into one dataset.

### Sample size selection

After we collected the data from the first animal (monkey A), we estimated a sample size that was required for a statistical test (*t*-test) between sham and each cooling condition to have a power of (1 – β) = 0.90 with a significance level of α = 0.01. This calculation was done with the SAMPSIZEPWR function of MATLAB. The mean and standard deviation of the null hypothesis was set to those of the sham condition and the sample size was estimated for endpoint error, which showed significant impairment in both PMd and A5 cooling (Fig. 3c). Results showed that the optimal sample size was 9 and 11 sessions for PMd and A5 cooling. Given the high variability of behaviours between animals, we set a minimal sample size for the second animal to 10 sessions for each condition.

### EMG analyses

In some recording sessions (n = 33 and 58 in monkey A and R, respectively), electromyographic (EMG) activity was recorded from upper-limb muscles by attaching surface EMG electrodes over each muscle belly (Delsys, Natwick MA, USA). Muscles were selected that predominantly contributed to flexion and extension movements at the shoulder and elbow (biceps, brachioradialis, brachialis, long/lateral triceps, anterior/middle/posterior deltoid, pectoralis major)^33^. Muscle activity was recorded at 4000 Hz, band-bass filtered (25-350Hz, 6th order Butterworth), full-wave rectified and downsampled to 1000 Hz before analysis.

EMG signals were aligned to perturbation onset and averaged across trials. Preferred torque direction (PTD) of each EMG was determined as the torque combination (SE+EF or SF+EE) that produced the larger response in a time window from 50 to 100 ms after perturbation onset. We identified EMG as perturbation responsive if the EMG response (50 – 100 ms after perturbation) in the PTD was significantly higher than that in an unloaded catch condition (paired *t*-test, *p* < 0.05). In total, we identified 293 EMG samples (n = 165 and 128 in monkey A and R) to be perturbation responsive and analyzed further (n = 140, 90 and 63 in sham, PMd and A5 cooling conditions, respectively).

EMG traces were first normalized by their mean activity during the last 2 sec of the trials when the monkey was countering the load in the PTD. This normalization value was calculated with data only from the pre-cool epoch, and then the same value was applied to all EMG data in the pre-cool, cool and post-cool epochs. Then, we averaged the normalized EMG traces across muscles separately for each cooling condition. From our pilot observation, we found that EMG signals had much higher noise than kinematic signals. Therefore, we performed a selection process of the EMG data based on the behavioural effects of cooling. Our behavioural analyses showed both PMd and A5 cooling increased endpoint errors (Fig. 3c). Therefore, we selected EMG data in sessions when the endpoint error was higher than the 90th percentile of the endpoint error from sham conditions. As a result, we selected 64 out of 153 EMG datasets (33 / 90 and 31 / 63 for PMd and A5 cooling). Importantly, we only used endpoint error for the selection and we used the same criteria for PMd cooling and A5 cooling. For the sham cooling data, no selection was applied (n = 140).

Muscle activity was compared across predefined epochs (baseline, 100-0 ms before perturbation onset; R1, 20-50 ms post-perturbation; R2, 50-75 ms post-perturbation; R3, 75-120 ms post-perturbation; voluntary 120-180 ms post-perturbation)^19,34^. Welch’s *t*-test was used to evaluate whether the binned muscle activity was significantly modulated from sham condition (*p* < 0.05 with Bonferroni correction).

### Temperature measurement in M1

In order to evaluate the change of M1 temperature during PMd cooling sessions, we recorded intracortical temperature of M1 on PMd cooling (n = 2) or dual cooling of PMd and A5 (n = 1) sessions. Prior to the temperature recording, we mapped the arm area of M1 using intracortical microstimulation (11 pulses, 333 Hz, 0.2 ms pulse width, ≤ 20μA). In each recording day, we inserted a thermocouple (HYP0-33-1-T-G-60-SMPW-M, Omega Engineering Inc, CT, USA) into the arm area of M1. We then started the experimental procedure to collect pre-cool, cool and post-cool epochs. The temperatures were sampled at 4000 Hz along with the kinematic signals.

Mean temperature during pre-cool (for 5 min before cooling onset) and cool (for 5 min before cooling offset) epochs were compared in M1 and PMd (probe temperature) separately (Fig. S3). Significant modulation of temperature was evaluated with paired *t*-test (*p* < 0.05 with Bonferroni correction)

### Model simulation

We used an optimal feedback control (OFC) model developed in our previous report^35^. We considered the translation of a single-point mass (*m* = 1 kg) in one dimension. The control system was described by the following differential equations:

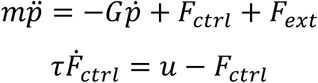

where *m* is mass, *p*(*t*) is the position of the point mass as a function of time (*t*), *G* is the viscous constant, *F*_*ctrl*_ and *F*_*ext*_ are the control and external forces, respectively, *u* is the motor command and *τ* is the time constant of the linear filter for the motor command. Each trial began with *F*_*ext*_ set to 0 Nm (no external load) and after 500ms *F*_*ext*_ was suddenly changed to +2 Nm and maintained for another 1000 ms until the end of the trial (a step-torque perturbation). *G* was set to 1 N·s·m^−1^, and *τ* was set to 40 ms, which is compatible with the first approximation of muscle dynamics^36^. Stochastic dynamics and noise disturbances are described in a discrete time system with a 10-ms time step:

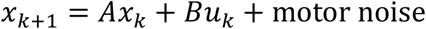

where *x*_*k*+*1*_ is the state vector at time *k*+*1*, *A* and *B* are matrices that describe the system dynamics, and *u*_*k*_ is the motor command at time *k*. Motor noise is a signal dependent Gaussian noise with variance of (0.125)^2^ × *u*^37^. The state vector is represented with the four-dimensional vector,

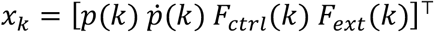

which was augmented with previous states to take feedback delays into account^38^. The feedback delay was set to 50ms (5 time steps) to reflect the transmission delay of the long-latency response^17^. This resulted in the augmented state vector with 24 dimensions.

The feedback signal at each time step (*y*_*k*_) can be written as,

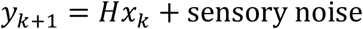

where *H* is the observation matrix, which allows the system to observe only the most delayed state (50ms before). Otherwise, the system is fully observable. Sensory noise is an additive Gaussian noise with variance 10^−10^. To compensate the time delay of the feedback signal, the OFC model includes an optimal linear state estimator (Kalman filter) that consists in weighting prior beliefs about the next state of the system with sensory feedback to derive a maximum likelihood estimate of the system state. Let 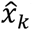 be the estimated state at time *k*. The prior belief about the next state (*x*\*_*k*+*1*_) is defined as

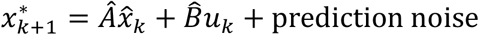

where 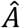 and 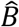 are the internal model of the system dynamics, *A* and *B*. Prediction noise is an additive Gaussian noise with variance 10^−8^. Then, the feedback correction yields the state estimation by taking *y*_*k*+*1*_ into account as

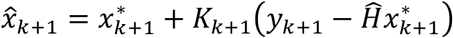

where *K*_*k*+*1*_ is the Kalman gain at time *k*+*1* and 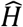 is the internal model of the observation matrix, *H*. Finally, the feedback control policy is defined as

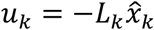

where *L*_*k*_ represents the optimal feedback gain at time *k*.

The cost function for the task was defined as:

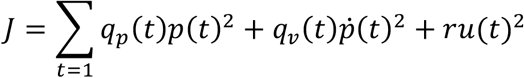

where *J* is the total-cost including error cost (position and velocity) and motor cost over a time-course of the trial. *q*_*p*_ and *q*_*v*_ are time-dependent factors which define the cost of position and velocity errors, respectively. Both were set to 1 before and 500 ms after the perturbation, but they were set to 0 between 0 and 500 ms after the perturbation. This means that the model is required to stay at (*p, ṗ*) = (0, 0) except for just after the perturbation. *r* is a constant factor which defines the cost of control and it was set to 10^−6^.

With these definitions of the system dynamics, feedback signals, noise parameters, and cost functions, we computed the optimal feedback gains (*L*) and Kalman gains (*K*) following algorithms adapted for the presence of signal-dependent noise^38,39^.

### Simulation for cortical deactivations

To apply deactivation of the optimized model parameters, we used two different methods: downscaling and noise addition. For downscaling, we multiply a scalar between 0 – 1 to deactivate the parameter. For example, when we deactivate a parameter by 20%, we multiply 0.8 (= 1 – 0.2) to the optimized value of the parameter. The rational of this method is that cortical cooling is known to reduce neural excitability at mainly post-synaptic terminals^40^, suggesting that cooling reduced the gain to the pre-synaptic inputs. Another possibility is that cortical cooling adds some noise to the neural computations. To replicate this scenario, we added scaled Gaussian noise to the target parameters. To control for deactivation size, the noise was chosen from a normal distribution whose standard deviation was scaled with the absolute value of each parameter:

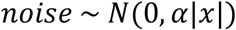

where |*x*| is absolute of the optimal value for the target parameter and *α* is a scaling factor of added noise chosen from 0 to 1. This means that noise size was normalized to the coefficient of variation (CV): *α* = 0 indicates that no noise was added (0% CV), whereas *α* = 1 indicates that noise chosen from normal distribution with standard deviation of |*x*| (100% CV).

With either deactivation method, we applied deactivation to each target parameter and evaluated the behavioural measures with the same algorithms used for the monkey behavioural analyses (Fig. 2d). Then we applied a two-way ANOVA (deactivation size × deactivated parameters) and *post hoc* paired *t*-tests to evaluate whether the behavioural measures were modulated from the optimal conditions (*p* < 0.05 with Bonferroni correction). Deactivation size for downscaling was chosen from 0 – 100% with a 5% step size, whereas deactivation size for noise addition was chosen from 0 – 40% with a 2 % step size.

## SUPPLEMENTARY MATERIALS

**Figure S1.**
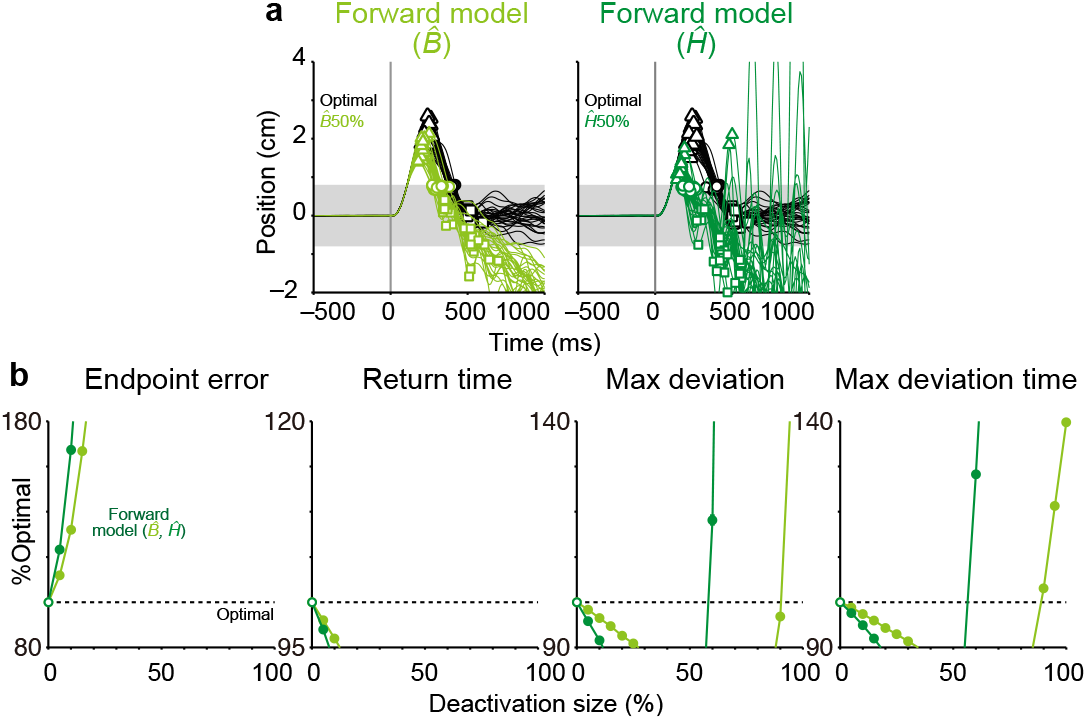
Deactivation of internal model parameters (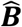 and 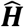). **a**, Response of the model in deactivated conditions. Same format as Fig. 1c but for 50% deactivation of 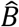 (light green) and 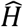 (green). **b**, Performance measures impaired by the deactivation of model parameters. Same format as Fig. 1d (filled circles, *t*-test, *p* < 0.05).

**Figure S2.**
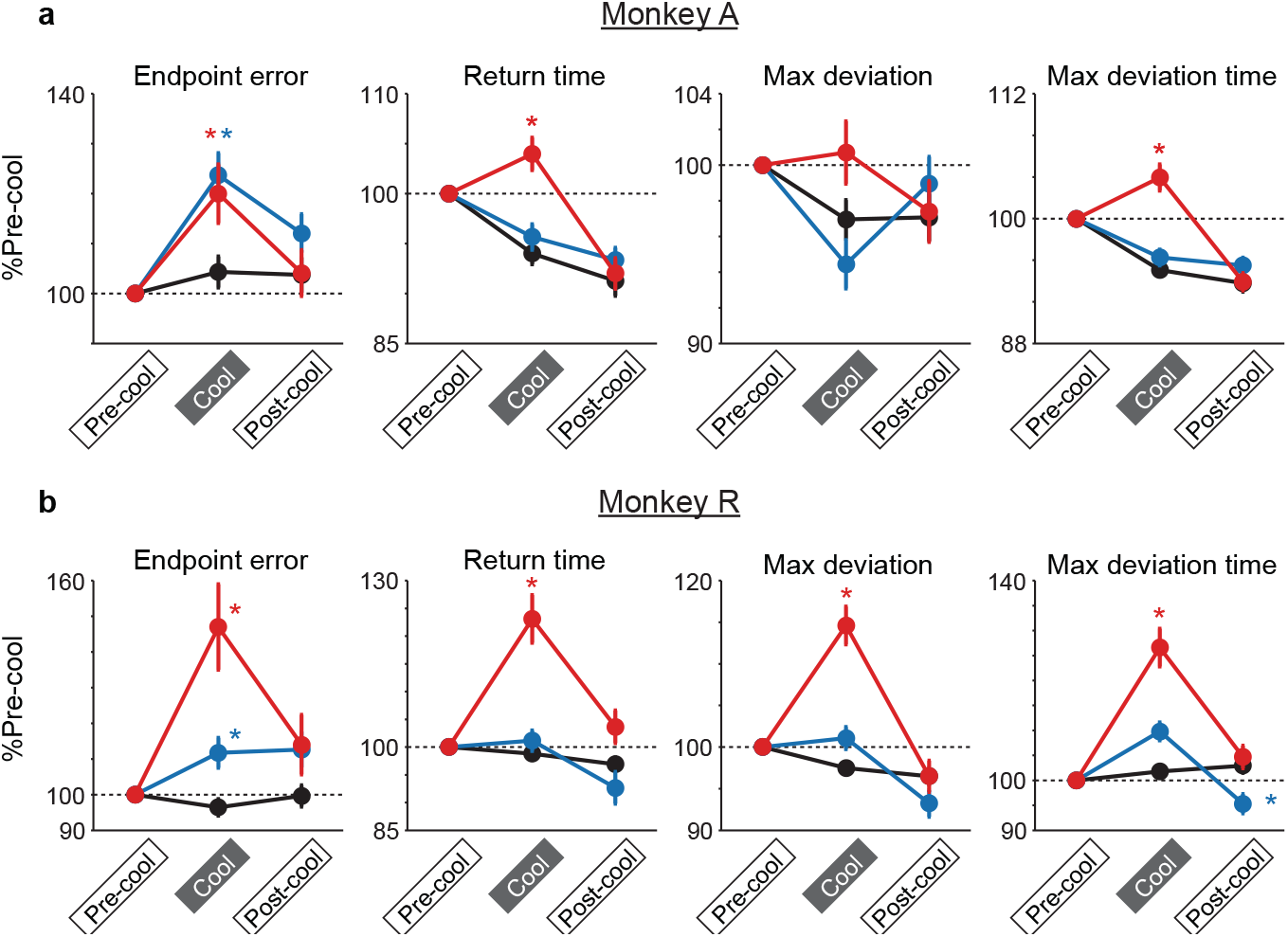
Behavioural results for each monkey (monkey A or R). Performance measures for PMd (red), A5 (blue) and sham (black) cooling sessions for monkey A (**a**) and monkey R (**b**). Same format as Fig. 3c. Error bars, SEM. * Significant difference from sham condition (*t*-test, *p* < 0.05). The results confirmed that our main finding holds across animals: PMd cooling impaired spatial accuracy (endpoint errors) and response speed (return time, max deviation, max deviation time), whereas A5 cooling impaired spatial accuracy but not response speed.

**Figure S3.**
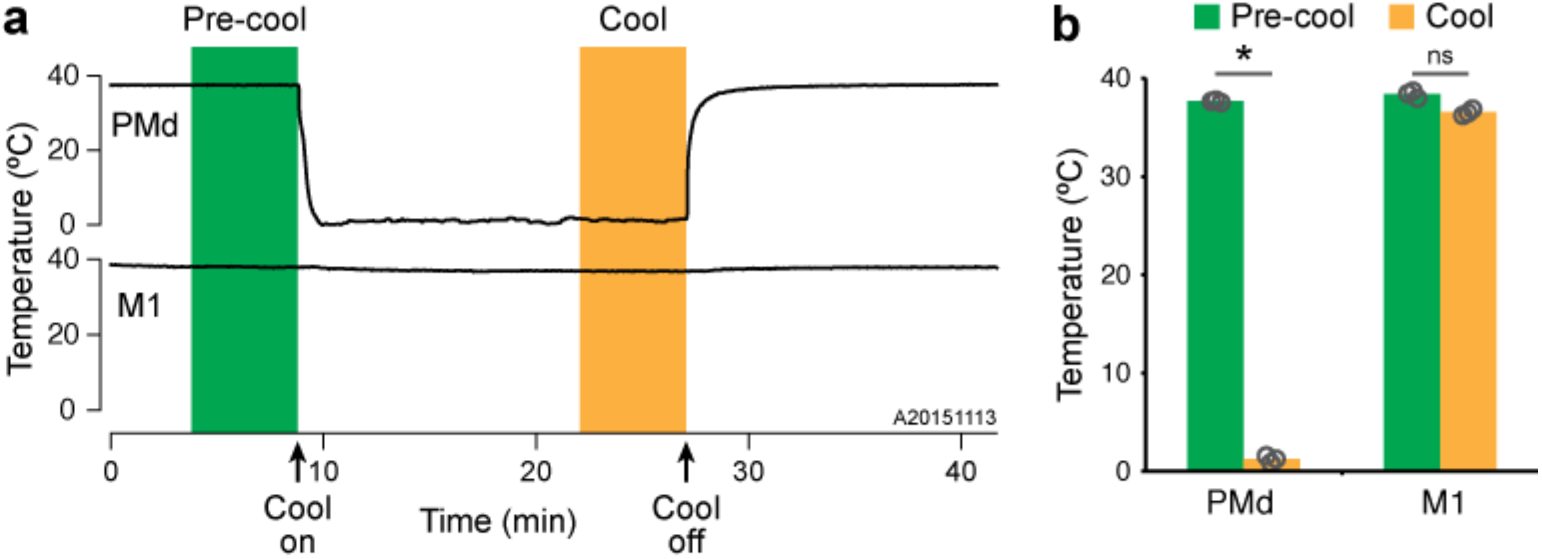
PMd and M1 temperature changes during PMd cooling. **a**, Example of cortical temperature measured from M1 during a PMd cooling experiment. Cooling probe on PMd was cooled to 1 °C and the probe temperature was measured (top). An independent thermocouple was acutely inserted to M1 arm area (bottom). **b**, Averaged temperature of PMd probe and M1 in pre-cool and cool epochs of PMd only (n = 2) and dual PMd and A5 cooling (n=1). * Significant difference between pre-cool and cool conditions (*t*-test, *p* < 0.05). ns, non-significant. Note that during PMd cooling or dual cooling, a change of M1 temperature was minimal (< 2°C) and did not reach significant limit.

**Figure S4.**
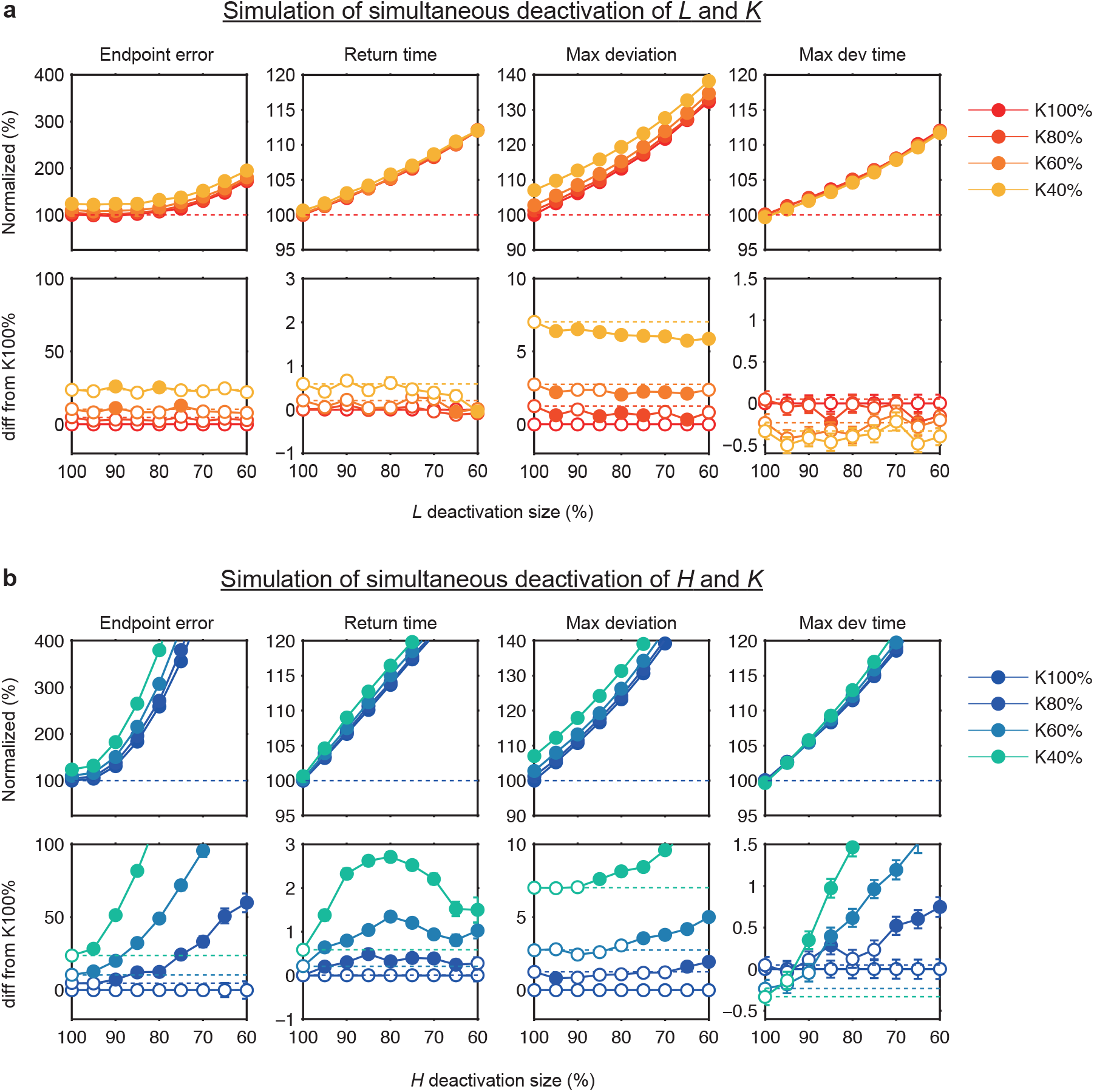
Simulation of the effect of dual deactivations. **a**, **b**, Results for dual deactivation of feedback gain and Kalman gain (*L* & *K*, **a**) and observation matrix and Kalman gain (*H* & *K*, **b**). Top: Performance measures were normalized to the optimal condition (dashed line). Bottom: Difference from the *K*100% condition. Horizontal dashed lines denote the linear sum of the deactivation of parameters (*L* & *K* or *H* & *K*). Filled circle, significant difference from the linear sum (*t*-test, *p* < 0.05). Note that the effects are linear or sublinear for *L* & *K* deactivation, whereas they are supralinear for *H* & *K*.

**Figure S5.**
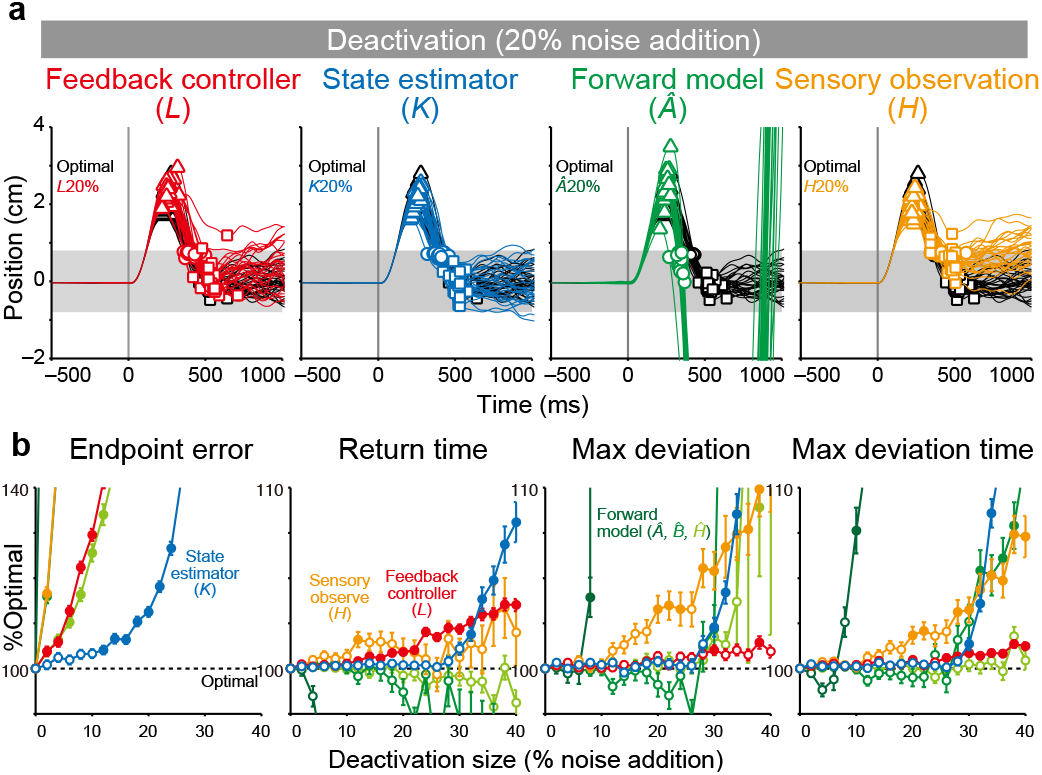
Simulation of cortical deactivation with noise addition. **a**, Response of the model when a scaled Gaussian noise (20% coefficient of variation) was added to each parameter. Same format as Fig. 1c. **b**, Impairment of behavioural measures as a function of deactivation size. Filled circles, significant difference from optimal condition (*t*-test, *p* < 0.05). Note that this deactivation method produced largely similar effects to the downscaling (Fig. 1). First, the deactivation of feedback gain (*L*) impaired both spatial accuracy (endpoint error) and response speed (return time), whereas the deactivation of Kalman gain (*K*) impaired the endpoint error but less effect on response speed at smaller deactivation size (10 – 30 % coefficient of variation). Second, the deactivation of forward models 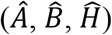 led severe oscillations. Finally, deactivation of sensory observation (*H*) impaired both spatial accuracy and response speed (max deviation and max deviation time).

### Supplementary Discussion: Simulation of cortical cooling with noise addition

In addition to downscaling, we also simulate the effect of cortical cooling by adding scaled Gaussian noise to each OFC parameter (see Methods). This method was based on an assumption that cortical cooling increases noise on neural processing. Results showed that the noise addition produced qualitatively similar effects on behavioural parameters to the downscaling, shown in Fig. 1. First, the deactivation of feedback gain (*L*) impaired both spatial accuracy (endpoint error) and response speed (return time), whereas the deactivation of Kalman gain (*K*) impaired the endpoint error but less for return time at smaller deactivations (10 – 30 % coefficient of variation). Second, the deactivation of forward models 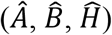 led to severe oscillations. Lastly, deactivation of sensory observation (*H*) impaired both spatial accuracy and response speed (max deviation and max deviation time).

It is noteworthy, however, that there were some discrepancies between simulation results with noise addition and the cortical cooling. First, the max deviation and max deviation time was less impaired by the deactivation of feedback gain (*L*) or Kalman gain (*K*). This is inconsistent with the results of PMd cooling, where all these parameters were significantly impaired (Fig. 3c). Second, a smaller deactivation of parameters was not able to predict the results of milder cooling of PMd (8 °C in probe temperature), which impaired response speed but not spatial accuracy. Although both mechanisms (reduction of gain and increase of noise) might underlie the effect of cortical cooling, the downscaling of model parameters appears to be a better model for the effects of cortical cooling.

